# Meiotic and mitotic aneuploidies drive arrest of in vitro fertilized human preimplantation embryos

**DOI:** 10.1101/2022.07.03.498614

**Authors:** Rajiv C. McCoy, Michael C. Summers, Abeo McCollin, Christian S. Ottolini, Kamal Ahuja, Alan H. Handyside

**Affiliations:** Department of Biology, Johns Hopkins University, 3400 N. Charles Street, Baltimore, Maryland, 21212, USA; London Women’s Clinic, 113-115 Harley Street, Marylebone, London, W1G 6AP, UK; School of Biosciences, University of Kent, Canterbury, Kent, CT2 7NJ, UK; London Women’s Clinic, The Chesterfield, Nuffield Health Clinic, 3 Clifton Hill, Bristol, BS8 1BN, UK; The Evewell, 61 Harley St, London, W1G 8QU, UK; Department of Maternal and Fetal Medicine, University College London, 86-96 Chenies Mews, London, WC1E 6HX UK

## Abstract

The high incidence of aneuploidy in early human development, arising either from errors in meiosis or postzygotic mitosis, is the primary cause of pregnancy loss, miscarriage, and still birth following natural conception as well as *in vitro* fertilization (IVF). Preimplantation genetic testing for aneuploidy (PGT-A) has confirmed the prevalence of meiotic and mitotic aneuploidies among blastocyst-stage IVF embryos that are candidates for transfer. However, only about half of normally fertilized embryos develop to the blastocyst stage *in vitro*, while the others arrest at cleavage to late morula or early blastocyst stages. To achieve a more complete view of the impacts of aneuploidy, we applied a validated method of PGT-A to a large series (n = 909) of arrested embryos and trophectoderm biopsies. We then correlated observed aneuploidies with abnormalities of the first two cleavage divisions using time lapse imaging (n = 843). The combined incidence of meiotic and mitotic aneuploidies was strongly associated with blastocyst morphological grading, with the proportion ranging from 20% to 90% for the highest to lowest grades, respectively. In contrast, the incidence of aneuploidy among arrested embryos was exceptionally high (94%), dominated by mitotic aneuploidies affecting multiple chromosomes. In turn, these mitotic aneuploidies were strongly associated with abnormal cleavage divisions, such that 51% of abnormally dividing embryos possessed mitotic aneuploidies compared to only 23% of normally dividing embryos. We conclude that the combination of meiotic and mitotic aneuploidies drives arrest of human embryos *in vitro*, as development increasingly relies on embryonic gene expression at the blastocyst stage.

## Introduction

Following natural conception, many human embryos are chromosomally abnormal and are progressively eliminated through preclinical pregnancy loss, miscarriage, and still birth, such that the overall incidence of these abnormalities detected in newborns is less than 1% (Gardner and Amor, 2018; Moorthie *et al.*, 2018). Observed chromosome abnormalities include genome-wide abnormalities in ploidy (e.g., triploidy), as well as whole and segmental aneuploidies of individual chromosomes (Levy *et al.*, 2014). Whole chromosome aneuploidy frequently arises through errors in meiosis, predominantly in females, resulting in aneuploid oocytes. Risk of such maternal meiotic aneuploidies, particularly of the smaller and acrocentric chromosomes, increases exponentially for women over the age of 35 years in parallel with increasing risk of miscarriage (Nagaoka *et al.*, 2012). A similar pattern of aneuploidy is also observed following *in vitro* fertilization (IVF) (Segawa *et al.*, 2017; Ishihara *et al.*, 2020). Hence, preimplantation genetic testing for aneuploidy (PGT-A) at the blastocyst stage, by trophectoderm biopsy and next-generation sequencing (NGS)-based chromosome copy number analysis, is now widely used for the selection of viable euploid blastocysts for transfer (Rosenwaks and Handyside, 2018).

Unlike previous methods used for PGT-A, including, for example, array comparative genomic hybridization (aCGH), NGS exhibits a linear relationship between normalized read depth and chromosome copy number, while also offering superior resolution for detecting segmental abnormalities (Fiorentino *et al.*, 2014). With multiple trophectoderm cells (typically 3-10 cells) biopsied at the blastocyst stage, this has enabled the identification of both whole and segmental chromosome aneuploidies with copy numbers ranging from those expected for trisomies or monosomies (full changes) to low or intermediate copy number changes (McCoy, 2017). By identifying meiotic errors in polar bodies and trophectoderm biopsies using SNP genotyping and karyomapping in parallel with NGS-based PGT-A, we recently demonstrated that, with one exception, all female meiotic aneuploidies resulted in copy number changes exceeding 70% of full changes in the corresponding trophectoderm biopsies or whole arrested embryos (Handyside *et al.*, 2021). In contrast, most non-meiotic (presumed mitotic origin) aneuploidies had copy number changes ranging from 30-70% although a minority exceeded the 70% threshold and may have resulted from chromosome missegregation in the first mitotic cleavage divisions.

Despite improvements in embryo culture, only about half of normally fertilized embryos reach the blastocyst stage, while the remainder arrest at various cleavage, late morula, or early blastocyst stages (Biggers and Summers, 2008; Summers *et al.*, 2013; Sfontouris *et al.*, 2016; Werner *et al.*, 2016). As early as 1993, Munné and colleagues demonstrated the association between aneuploidy and asymmetric cleavage and embryo arrest using multicolour interphase fluorescent in situ hybridisation with chromosome specific probes for X, Y and 18 (Munné *et al.*, 1993). Since then, with the introduction of comprehensive chromosome testing methods, numerous studies have confirmed this association (Fragouli *et al.*, 2013; Capalbo *et al.*, 2014; Qi *et al.*, 2014; Maurer *et al.*, 2015). However, all of these studies were limited by the use of earlier methodologies that did not discriminate between full and intermediate copy number changes for all chromosomes, limited sampling of embryo cells, or analysis of selected clinical-grade embryos only (reviewed by Regin *et al.*, 2022).

Here, we use the established copy number thresholds (Handyside et al., 2021) to discriminate meiotic- and mitotic-origin aneuploidies based on NGS of a large sample of arrested embryos and trophectoderm biopsies of blastocysts irrespective of their morphological grade. By testing both arrested embryos and blastocysts from the same IVF cycles, we infer the relative contributions of various forms of aneuploidy to preimplantation embryo arrest. Furthermore, as morphokinetic parameters are reported to be altered in aneuploid embryos (Campbell *et al.*, 2013; Vera-Rodriguez *et al.*, 2015; Lagalla *et al.*, 2017), time lapse analysis was used to identify abnormalities particularly in the first cleavage divisions, and correlate these with aneuploidy and developmental outcomes. Finally, by mitigating the survivorship biases that affect most retrospective studies, our study refines estimates of the incidences of meiotic and mitotic aneuploidy and their relationship with maternal age. Together our study offers a detailed view of chromosome and cleavage abnormalities in human preimplantation embryos and their contributions to embryonic mortality.

## Results

Between January 2016 and June 2018, 125 patients (mean 38.9 years at oocyte retrieval; range 30-45 years) underwent a total of 165 IVF cycles with extended embryo culture to the blastocyst stage and biopsy of 5-10 trophectoderm cells on days 5-7 postinsemination for PGT-A. Excluding an additional 22 cycles in which all embryos arrested and were not tested by PGT-A (a total of 52 zygotes with two pronuclei [2PN]), an average of 3.8 of 7.2 (53%) embryos reached the blastocyst stage per cycle—a proportion that declined with maternal age (Quasibinomial GLM: *β* = -0.080, SE = 0.022, *p* = 3.7 x 10^-4^; Table S1). Considering all 1232 normally fertilized (2PN) zygotes and excluding possible triploid embryos subsequently identified by PGT-A (see Methods), 622 (50.5%) embryos developed to the blastocyst stage and 610 (49.5%) embryos arrested between the zygote and late morula/early blastocyst stages (Figure 1). A total of 909 (73.4%) embryos derived from 2PN zygotes were tested with PGT-A, including 612 of the 622 (98.4%) blastocysts and 297 of the 610 (48.7%) arrested embryos. Notably, this includes 85 (51.5%) cycles in which all embryos were tested, as well as 80 cycles (48.5%) in which only a subset (mean = 24.1%) of arrested embryos were tested.

**Figure 1.**
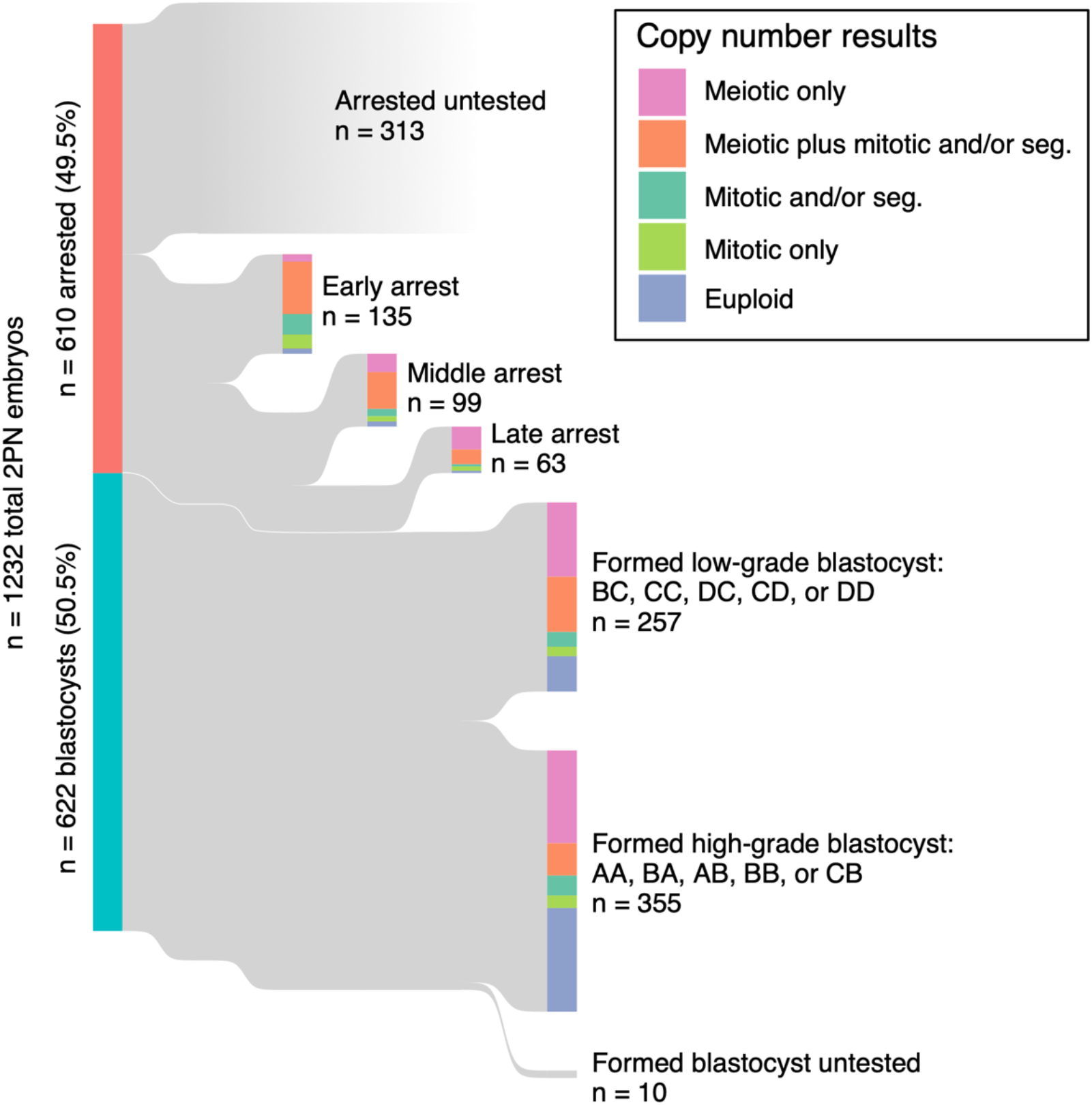
Developmental outcomes of 1232 normally fertilized (2PN) embryos and their associated PGT-A results. A total of 909 (73.8%) of embryos were tested with PGT-A, including 297 of 610 (48.7%) arrested embryos and 612 of 622 (98.4%) blastocysts. Early arrest: cleavage stages; ≤ 10 cells. Mid arrest: >10 cells, but pre-compact morula. Late arrest: compact morula to cavitating (non-expanded) blastocyst.

While we opted to analyze all 165 cycles to maximize statistical power, we also replicate key results in the subset of 85 cycles where all embryos were tested, as presented in the supplementary materials. We highlight the investigation of arrested embryos as well as poorer quality blastocysts (257 embryos with morphological grades BC, CC, CD, DC, or DD) as important aspects of our study, as such embryos are not as frequently tested in clinical settings because they are not typically considered as good candidates for either biopsy, vitrification, or transfer.

### Meiotic and mitotic aneuploidies are prevalent among preimplantation embryos

Across 909 tested embryos, 206 (22.6%) were euploid while 703 (77.3%) possessed whole or segmental aneuploidies of one or more chromosomes. To gain insight into the origins of aneuploidies and their consequences for development, we distinguished putative meiotic and mitotic aneuploidies based on their PGT-A copy number profiles (see Methods). Along with 50 aneuploidies of entire sex chromosomes, for which discerning mosaic aneuploidy poses unique challenges (see Methods), we discovered 1154 putative meiotic and 1051 putative mitotic aneuploidies of whole autosomes. We additionally identified 358 large segmental aneuploidies (on the scale of entire chromosome arms), including 351 affecting autosomes and 7 affecting sex chromosomes.

Consistent with previous work (e.g., McCoy *et al.*, 2015b), the meiotic aneuploidies disproportionately impacted chromosomes 15, 16, 19, 21, and 22, whereas mitotic aneuploidies exhibited similar frequencies across all autosomes (Figure 2A; Figure S1A). Moreover, we confirmed that the putative meiotic aneuploidies possessed a strong association with maternal age (Quasibinomial generalized linear model [GLM]: *β* = 0.261, standard error [SE] = 0.032, *p* value [*p*] = 1.68 x 10^-12^; Figure 3A), whereas putative mitotic aneuploidies exhibited no significant age association (Quasibinomial GLM: *β* = 0.009, SE = 0.033, *p* = 0.799; Figure 3B).

**Figure 2.**
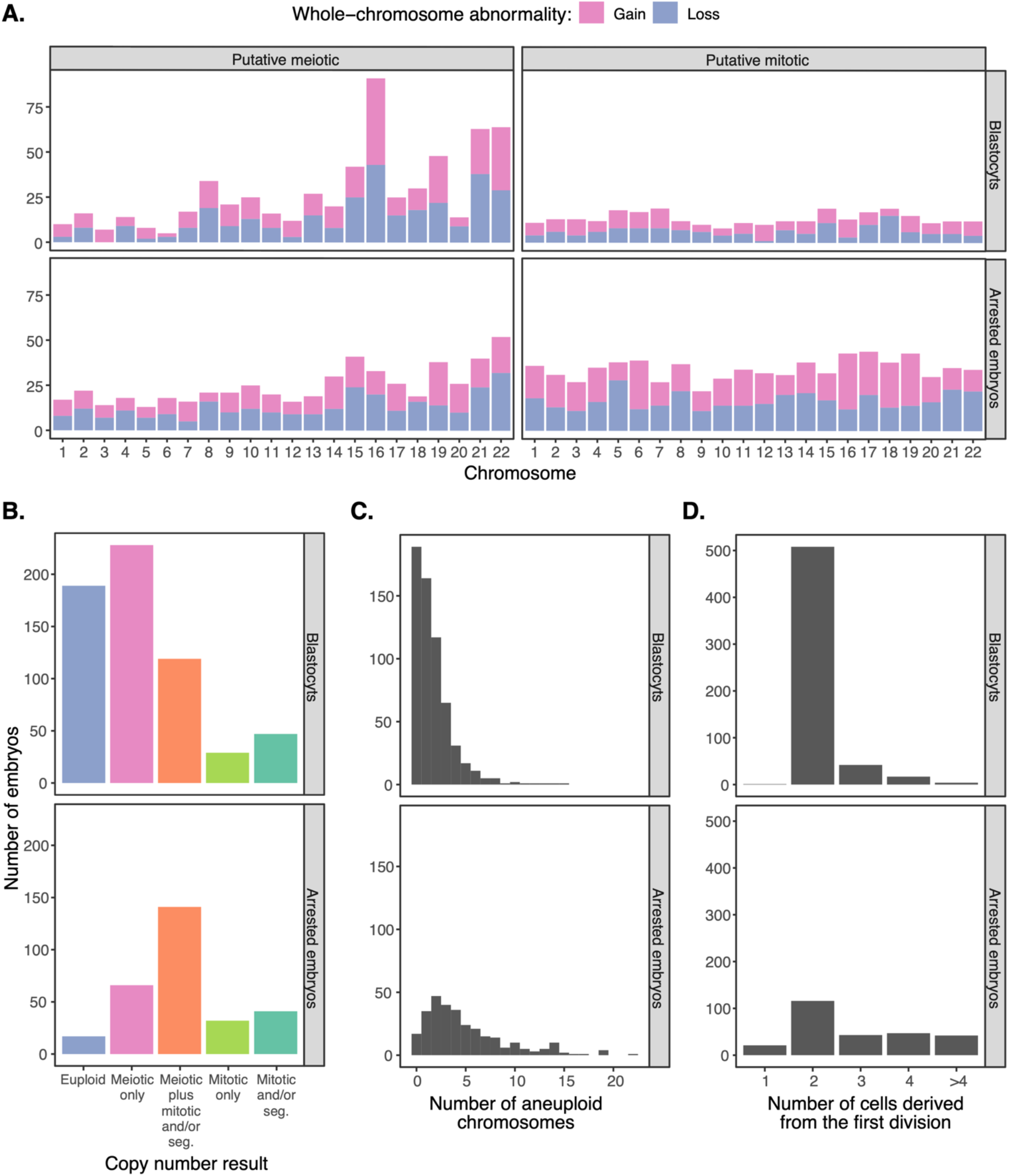
Characteristics of blastocysts and arrested embryos. **A.** Chromosome-specific counts of putative meiotic (i.e., full copy number change) and putative mitotic (i.e., intermediate copy number change) whole-chromosome gains and losses observed in blastocysts and arrested embryos as determined with PGT-A. Only autosomes are depicted, as distinguishing meiotic and mitotic origins of aneuploidies affecting sex chromosomes poses unique challenges (see Methods). **B.** Counts of arrested embryos versus developing blastocysts, stratified by PGT-A copy number result category. **C.** Counts of arrested embryos versus developing blastocysts, stratified by total number of aneuploid chromosomes. **D.** Counts of arrested embryos versus developing blastocysts, stratified by number of cells present after the first mitotic division.

**Figure 3.**
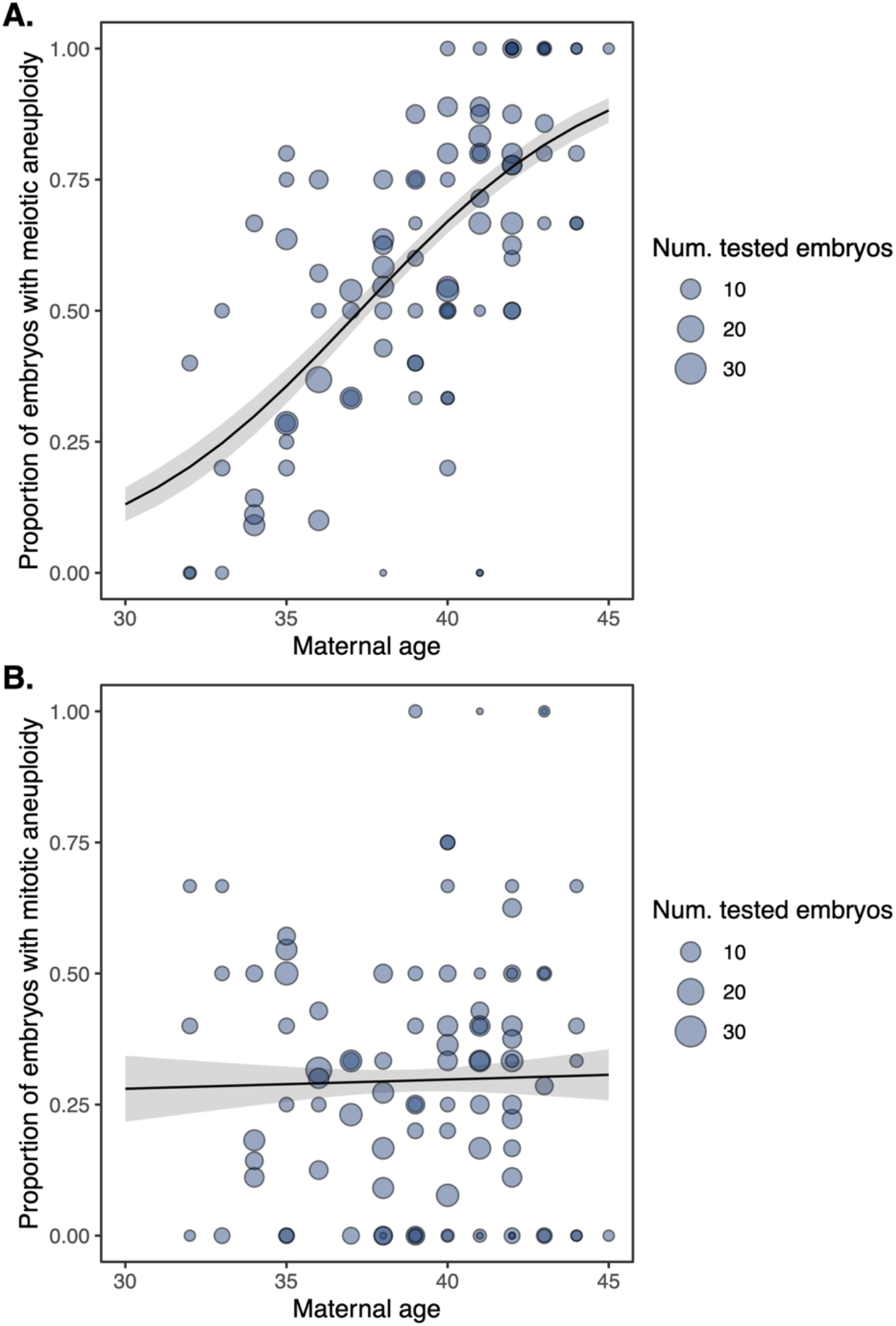
Observed rates of meiotic (**A.**) and mitotic (**B.**) aneuploidy in relation to maternal age, as measured from a large sample of arrested embryos and blastocysts. Each data point represents a distinct IVF case. Lines represent predictions from binomial generalized linear models fit to the data, with standard errors of the predictions indicated in gray.

We observed no significant co-occurrence of meiotic and mitotic aneuploidies affecting different chromosomes of the same embryos (Fisher’s Exact Test: odds ratio [OR] = 1.32, 95% confidence interval [CI; 0.99, 1.77], *p* = 0.055), suggesting that these error mechanisms are largely independent. However, when restricting to the 612 blastocyst-stage embryos, we observe a slight enrichment of mitotic aneuploidy among embryos already affected by meiotic aneuploidy (Fisher’s Exact Test: OR = 1.54, 95% CI [1.02, 2.36], *p* = 0.036). While the wide confidence intervals suggest that our power for detecting such an effect is limited in both cases, the latter observation raises the intriguing hypothesis that following embryonic genome activation, the functional effects of meiotic aneuploidies may include impacts on the mitotic machinery, compromising the fidelity of chromosome segregation.

### Mitotic aneuploidies disproportionately contribute to preimplantation arrest

The overall incidence of aneuploidy among arrested embryos was 94%, compared to 69% in tested blastocysts. Both putative meiotic and putative mitotic whole-chromosome aneuploidies were enriched among arrested embryos compared to embryos that developed to the blastocyst stage, though the effect was much stronger for mitotic aneuploidies (Fisher’s Exact Test: OR = 6.02, 95% CI [4.40, 8.28], *p* = 4.22 x 10^-33^) compared to meiotic aneuploidies (Fisher’s Exact Test: OR = 1.62, 95% CI [1.20, 2.20], *p* = 0.0012).

For the purpose of downstream analysis and visualization, we further grouped embryos into one of five mutually exclusive categories: (1) euploid, (2) meiotic whole-chromosome aneuploidy only, (3) meiotic whole-chromosome aneuploidy in combination with mitotic whole-chromosome aneuploidy or segmental aneuploidy (4) mitotic whole-chromosome aneuploidy only, and (5) segmental aneuploidy with or without mitotic whole-chromosome aneuploidy (Table 1). The distribution of embryos among these categories was significantly different for arrested embryos compared to trophectoderm biopsies of developing blastocysts (Figure 2B; Pearson’s Chi-squared Test: *X^2^* [4, N = 909] = 143.4, *p* = 5.41 x 10^-30^; Figure S1B).

**Table 1.** Incidence of various forms of aneuploidy in whole arrested embryos (see Methods for description of arrested embryos) versus trophectoderm biopsies of developing blastocysts as determined by preimplantation genetic testing for aneuploidy (PGT-A). Our approach for distinguishing putative meiotic and mitotic aneuploidies based on chromosome copy number results is detailed in the main text.

| Putative chromosome status | Abbreviation | Whole arrested embryos | Blastocyst-stage embryos (trophectoderm biopsies) |
| --- | --- | --- | --- |
| Euploid | Euploid | 17 (6%) | 189 (31%) |
| Meiotic whole-chromosome aneuploidy only | Meiotic only | 66 (22%) | 228 (37%) |
| Meiotic whole-chromosome aneuploidy with mitotic whole-chromosome aneuploidy or segmental aneuploidy | Meiotic plus mitotic and/or seg. | 141 (47%) | 119 (19%) |
| Mitotic whole-chromosome aneuploidy only | Mitotic only | 32 (11%) | 29 (5%) |
| Segmental aneuploidy with or without mitotic whole-chromosome aneuploidy | Mitotic and/or seg. | 41 (14%) | 47 (8%) |
| **Total** |  | 297 | 612 |

Whereas euploid embryos exhibited a 16% frequency of arrest (95% CI [10%, 25%]), embryos with solely meiotic aneuploidy (“Meiotic only”) arrested at a 36% frequency (95% CI [28%, 45%]), while embryos with mitotic aneuploidies (“Meiotic plus mitotic and/or segmental”, “Mitotic only”, “Mitotic and/or segmental”) arrested at much higher frequencies (ranging from 55% to 64%; Figure 4A,B; Figure S2A,B). Moreover, the number of aneuploid chromosomes per embryo was strongly associated with the frequency of arrest, with complex aneuploidies arresting in much higher proportions (Binomial GLMM: average marginal effect [AME] = 0.440, SE = 0.054, *p* = 4.04 x 10^-16^; Figure 2C; Figure 4C,D; Figure S1C; Figure S2C,D), building upon previous observations from smaller samples (Vega et al., 2014). Interestingly, maternal age was not associated with the frequency of embryo arrest after accounting for aneuploidy status of those embryos (Likelihood ratio test statistic = 0.57, *p* = 0.452), suggesting that the maternal age effect on IVF embryo loss is nearly exclusively mediated by aneuploidy.

**Figure 4.**
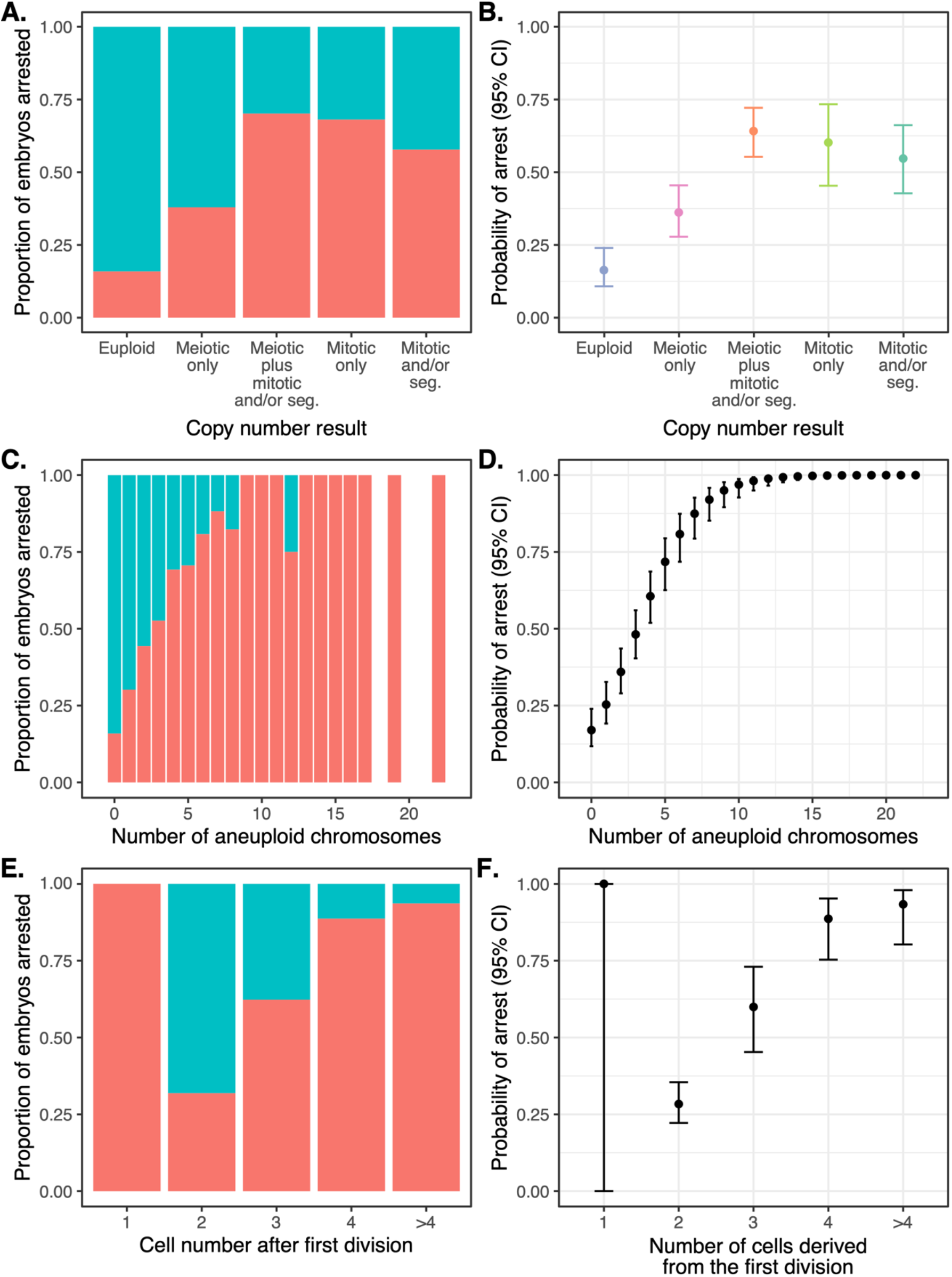
Data (left panels) and statistical modeling (right panels) of the proportion/probability of embryo arrest, stratifying on various patterns of chromosome copy number or cell division. Error bars denote 95% confidence intervals of estimates. **A.** Proportion of embryos arrested (red) versus unarrested (blue), stratifying on chromosome copy number pattern, as assessed by PGT-A. **B.** Statistical modeling of the data from panel A. **C.** Proportion of embryos arrested (red) versus unarrested (blue), stratifying on the number of aneuploid chromosomes. **D.** Statistical modeling of data from panel C. **E**. Proportion of embryos arrested (red) versus unarrested (blue), stratifying on the number of cells observed after the first mitotic division, where 2 is normal. **F.** Statistical modeling of data from panel E.

Given the strong association between aneuploidy and pregnancy loss, one quantity of fundamental interest is the proportion of embryos derived from euploid zygotes that arrest during preimplantation development, as these compose the initial pool of potentially viable zygotes whose success we seek to maximize. The ability to estimate this parameter has been hindered by the fact that early-arresting embryos are not typically tested in clinical settings. Within our data, the embryos derived from euploid zygotes include any embryo that lacks evidence of meiotic aneuploidy. Embryos in this group arrested at a frequency of 38% (95% CI [31%, 45%]), whereas embryos derived from the remaining aneuploid zygotes arrested at a frequency of 50% (95% CI [44%, 57%]).

### Abnormal cell divisions drive lethal mitotic aneuploidies

To gain insight into the relationship between abnormal early cell divisions and mitotic aneuploidies, we used time-lapse imaging (see Methods) to record the first two cell divisions of 843 embryos derived from 2PN zygotes following fertilization. Overall, 219 embryos (26.0%) exhibited an abnormal first division, while an additional 82 embryos (9.7%) exhibited an abnormal second division (Table S2). The former group includes 85 embryos (10.1%) that underwent abnormal division from one into three cells, largely due to precocious or multipolar cell divisions. We observed a strong association between abnormal cell division and putative mitotic aneuploidy, with 51% of abnormally dividing embryos possessing mitotic aneuploidies compared to only 23% of normally dividing embryos (Fisher’s Exact Test: OR = 3.41, 95% CI [2.50, 4.67], *p* = 6.93 x 10^-16^). No such relationship was observed between abnormal cell division and meiotic aneuploidy (Fisher’s Exact Test: OR = 1.20, 95% CI [0.89, 1.62], *p* = 0.217). As further expected, abnormal cell divisions were also strongly associated with embryonic arrest (Fisher’s Exact Test: OR = 10.86, 95% CI [7.67, 15.50], *p* = 2.72 x 10^-50^; Figure 2D; Figure S1D; Figure 4E,F; Figure S2E,F).

Consistent with this interpretation, the probability that embryos derived from euploid zygotes would arrest strongly depended on the outcomes of the first and second cell divisions. Specifically, the probability that an embryo derived from a euploid zygote would arrest was 12% (95% CI [7%, 19%]) when conditioning on normal first and second mitotic divisions compared to 34% (95% CI [27%, 42%]) for aneuploid zygotes. These proportions increase to 75% (95% CI [62%, 84%]) for euploid zygotes as well as 75% (95% CI [66%, 82%]) for aneuploid zygotes when conditioning on abnormal first or second cell divisions.

Notably, even embryos lacking meiotic or mitotic aneuploidies were much more likely to arrest if they experienced an abnormal first or second cell division (Fisher’s Exact Test: OR = 19.90, 95% CI [5.59, 90.50], *p* = 9.12 x 10^-8^). Though this accounts for only 13 embryos total (compared to 4 euploid embryos that arrested following normal cell divisions), the observed enrichment suggests that the fitness impacts of abnormal cell division are not solely attributable to aneuploidy.

### Aneuploidy is strongly associated with blastocyst morphology

Among embryos that survived to the blastocyst stage, we sought to understand the relationship between various forms of aneuploidy and blastocyst morphology, with the rationale that aneuploidies may compromise the organization and function of the differentiating cell lineages. Blastocysts were graded by assigning an ordinal letter grade (A through D) to the inner cell mass as well as the trophectoderm based on standardized morphological criteria (Table S3; Figure S3; Alpha/ESHRE, 2011). Notably, morphological grading was conducted at the time of embryo culture, prior to obtaining and thereby blind to the PGT-A results. Within our sample, the grades for the inner cell mass and trophectoderm were strongly correlated, consistent with the interpretation that ploidy status and other shared genetic and environmental factors simultaneously impact both cell types (Figure S4). We observed a strong association between aneuploidy status and blastocyst morphology, with aneuploidy rates ranging from 20% to 90% for embryos with the highest to lowest grades. Poor morphology embryos were significantly enriched for aneuploidy (Pearson’s Chi-squared Test: *X^2^* [9, N = 612] = 60.2, *p* = 1.23 x 10^-9^; Figure 5; Figure S5). This relationship was consistent for embryos affected by meiotic aneuploidies (Pearson’s Chi-squared Test: *X^2^* [9, N = 612] = 49.4, *p* = 1.38 x 10^-7^) as well as those affected with mitotic aneuploidies (Pearson’s Chi-squared Test: *X*^2^ [9, N = 612] = 49.3, *p* = 1.45 x 10^-7^) when compared to euploid embryos. Additionally, the presence of meiotic and/or mitotic aneuploidies was associated with later timing of blastocyst biopsy, indicating that aneuploidies tend to delay the process of blastocyst formation and expansion (LMM: AME = 0.220, SE = 0.052, *p* = 2.4 x 10^-5^; Figure S6).

**Figure 5.**
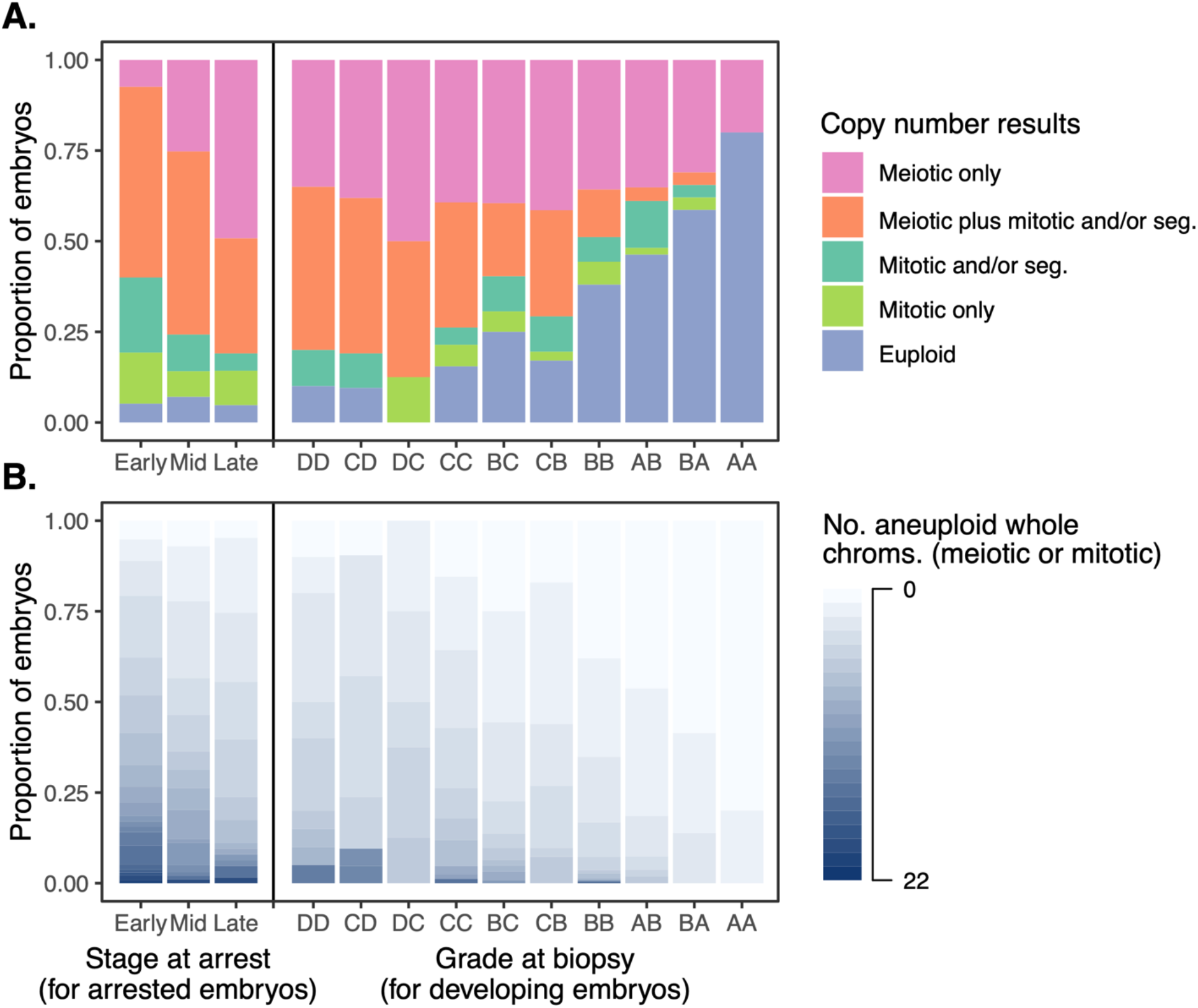
Chromosome copy number results (as assessed via PGT-A) across all tested embryos, stratifying by stage at arrest (see Methods for description of arrested embryos) or morphological grade (for embryos that formed blastocysts). ICM grade is listed first and TE grade is listed second. **A.** Copy number results assigned to categories, as described in Table 1. **B.** Copy number results depicted as counts of aneuploid chromosomes.

## Discussion

Here we applied a combination of genome-wide NGS-based copy number analysis and time-lapse imaging to investigate a large sample of both arrested and developing embryos, irrespective of their morphological grade or any other criteria, toward a more comprehensive view of the impacts of chromosome and cell division abnormalities on human preimplantation development. By comparing patterns of putative meiotic and mitotic aneuploidies across these samples and stages, using previously validated copy number thresholds (Handyside *et al.*, 2021), we observed that all categories of aneuploidy were enriched in arrested embryos versus developing blastocysts, reflecting their respective contributions to embryonic mortality. The leading cause of IVF embryo arrest appears to be lethal mitotic aneuploidies, which are in turn associated with abnormal first or second mitotic divisions and tend to affect multiple chromosomes simultaneously. Among surviving blastocysts, we further documented a strong association between patterns of aneuploidy and embryo morphology.

We note that the rates of aneuploidy observed in our data (77.3%) are higher than some previous studies (e.g., Franasiak *et al.*, 2014; Capalbo *et al.*, 2021). This observation is explained by the age distribution of the patient cohort, as well as the fact that most previous studies were based on retrospective analysis of data from embryos that were candidates for IVF transfer, systematically enriched for surviving embryos with good morphology and thus lower rates of both meiotic and mitotic aneuploidy (Figure 5A and 5B). By mitigating these biases, our data thus provide fundamental estimates of the underlying rates of meiotic and mitotic aneuploidy in human zygotes and early embryos (Figures 3A and 3B).

Studies in model systems have demonstrated that aneuploidy compromises cellular fitness (Torres *et al.*, 2007; Stingele *et al.*, 2012). Direct dosage effects of aneuploidy are known to exert proteotoxic and energy stress on cells, increased rates of mutation, and increased rates of chromosome mis-segregation, which may prove lethal during this critical early developmental transition. In particular, excess or depletion of proteins encoded on the aneuploid chromosomes confer stoichiometric imbalances among components of multiprotein complexes, inducing the formation of toxic protein aggregates (Zhu *et al.*, 2018; Brennan *et al.*, 2019; Li *et al.*, 2022). Recent work in aneuploid yeast suggests that these free proteins and protein aggregates increase the solute concentration within cells, causing chronic hypo-osmotic stress that impairs endocytosis (Tsai *et al.*, 2019).

The chromosome instability (CIN) and mitotic aneuploidy observed in the early cleavage-stage IVF human embryos may reflect defects in DNA repair, cell cycle progression, and intracellular signaling as a result of underlying imbalances in cell regulatory networks, metabolic flux, and cellular dynamics (Vanneste *et al.*, 2009; McCoy, 2017; Potapova and Gorbsky, 2017; Tšuiko et al., 2021; Sfakianoudis et al., 2021). For example, RNA sequencing results of aneuploid cells from early human embryos and blastocysts exhibit transcriptional profiles consistent with DNA damage (Vera-Rodriguez *et al.*, 2015; Licciardi *et al.*, 2019), changes and disruption of cell proliferation and the cell cycle (Starostik *et al.*, 2020; Maxwell *et al.*, 2022), p53 signaling, autophagy, and apoptosis (Groff *et al.*, 2019; Licciardi *et al.*, 2019; Sanchez-Ribas *et al.*, 2019; Maxwell *et al.*, 2022). These reports are consistent with a higher degree of chaotic aneuploidy observed in poorer quality and slower growing blastocysts (Figures 5A,B; Figures S5A,B; Figures S6A,B). Interestingly, recent work by Palmerola *et al.* (2022) confirmed a high level of incomplete DNA replication resulting in chromosome breaks and mitotic aneuploidy in human zygotes, referred to as replication stress. It is tempting to conclude that our findings are consistent with those of Palmerola *et al,* (2022) with the caveat that their studies involved the use of donated vitrified MII oocytes (see also Cavazza and Schuh, 2022).

Our data demonstrate that mitotic aneuploidies are prevalent and occur at similar frequencies for embryos derived from zygotes affected or unaffected with prior meiotic aneuploidies, in turn suggesting an intrinsic instability of cleavage-stage development and/or detrimental effects of embryo culture. It has been proposed, for example, that the blastomeres of early cleavage-stage human embryos function autonomously due to the absence of gap junctions and are particularly sensitive to environmental and metabolic perturbations (Brison *et al.*, 2014), perhaps explaining why, for example, a single blastomere only undergoes an abnormal unequal cleavage, such as a tripolar mitosis. By contrast, genome integrity at later stages of development may be more robust, following embryonic genome activation and the development of a transporting epithelium at the time of compaction.

It is perhaps worth noting that human embryo culture media provide only a partial representation (including a protein source, simple salts, energy substrates, and amino acids) of the natural environment to which embryos are exposed *in vivo* (Summers and Biggers, 2003). Consequently, stresses are invariably imposed on preimplantation embryos grown in chemically defined media. Human embryos appear robust and can develop in a wide selection of commercial media, indicating that they compensate and/or adapt to the imposed stresses (Biggers and Summers, 2008), but may ultimately succumb under extreme conditions (Puscheck *et al.*, 2015; Cagnone and Sirard, 2016). The precise environmental tipping points and genetic factors that influence the probability of human embryo arrest merit detailed investigation.

In their provocatively titled article, “Where have all the conceptions gone?”, Roberts and Lowe (1975) posited that embryonic mortality is the rule rather than the exception, occurring at a rate of ~80%. Highly cited reviews over subsequent decades offered qualitative support for this conclusion, estimating rates of 60-70% (Chard, 1991; Macklon *et al.*, 2002). Current evidence suggests that a high rate of preclinical losses, not failure of conception, is the main cause of low fecundity of humans, largely driven by chromosome abnormalities (Brosens *et al.*, 2022). By contrasting aneuploidies observed in arrested and unarrested embryos, our study supports this conclusion and clarifies the role of meiotic and mitotic aneuploidies in early embryonic mortality. Specifically, we show that severe mitotic errors, which frequently arise due to abnormalities in the initial postzygotic cleavage divisions, are the primary cause of mortality among IVF embryos.

Together, our results suggest that the transition from the cleavage to blastocyst stage may act as a strong bottleneck wherein numerous aneuploid embryos are eliminated prior to implantation. The timing of this bottleneck is consistent with the increasing reliance on embryonic gene expression as development proceeds through the cleavage, late morula, and early blastocyst stages (Tesarik *et al.*, 1987; Braude *et al.*, 1988; Yan *et al.*, 2013; Blakeley *et al.*, 2015).

## Material and Methods

### Study design and informed consent

This report represents a subset of patients that were part of a prospective cohort single center study of patients undergoing IVF with blastocyst vitrification-only and optional PGT-A (VeriSeqPGS, Illumina Inc) between January 2016 to June 2018, previously described by Gorodeckaja *et al.* (2019).

All procedures performed at the London Women’s Clinic, including quality control measures, were approved and licensed by the Human Fertilization and Embryology Authority (HFEA) in the United Kingdom in accordance with all relevant regulations and legislation. Details of ovarian stimulation protocols and laboratory procedures have been described in Gorodeckaja *et al.* (2019). No new procedures, protocols, or randomization were used in the program. Consequently, the study does not constitute human subjects research and accordingly approval from an Institutional Review Board was not required. Consenting of patients was carried out in accordance with the Code of Ethics of the World Medical Association (Helsinki Declaration). All patients undergoing PGT-A were consented by a qualified clinician and licensed genetic counselor. Written informed consent for genetic testing and follow up analysis, including the anonymous use of embryology data for statistical evaluation and research was reviewed, signed and witnessed. The Homewood Institutional Review Board of Johns Hopkins University determined that the analysis of these data does not qualify as federally regulated human subjects research (HIRB00011431).

### Time-lapse image analysis and blastocyst grading

Time-lapse incubation was used to maintain an uninterrupted embryo culture environment. Embryo development was monitored continuously up to seven days post insemination, or until blastocyst formation and expansion, if earlier, by time lapse imaging and software analysis (Geri® Connect, GeneaBiomedx, Sydney, Australia). Blastocysts were evaluated by assigning a letter grade (A through D) to the inner cell mass (ICM) and trophectoderm (TE) based on standardized morphological criteria (Table S3; Figure S1; Alpha/ESHRE, 2011). Grading of top quality blastocyst was more restrictive, whereas very poor quality blastocysts were given a D grade based on very few cells in the ICM and TE and evidence of cellular degeneration. Time lapse videos of all embryos were annotated daily using the manufacturer’s software and documentation in the patient’s laboratory record. Assessment included several time points in the development of each embryo: (1) formation of pronuclei, (2) first cleavage division, (3) second cleavage division, (5) cell compaction, (6) cavitation, (7) expanded blastocyst, or (8) embryo arrest image prior to processing embryos for PGT-A. The division pattern was recorded for each embryo as per Ottolini *et al*. (2017) and based on the number of cells after each division to indicate either normal (e.g., 1-2-4 cells) or abnormal (e.g., 1-3-6 cells) cleavage patterns (Rubio *et al.*, 2012; Ciray *et al.*, 2014; Kalatova *et al.*, 2015; Coticchio *et al.*, 2021). Categories of abnormal division included multipolar, precocious, reverse, and failed cleavage (McCollin *et al.*, 2020). “Multipolar” cleavage refers to the direct cleavage of the zygote (or a daughter cell) into three or more cells. “Precocious” cleavage refers to a rapid division pattern where the zygote (or a daughter cell) undergoes a normal 1→2 cell cleavage, followed by a subsequent premature division to produce 3 or more blastomeres. “Reverse” cleavage refers to the resorption of blastomeres after cytokinesis. “Failed” cleavage refers to multiple rounds of karyokinesis without cytokinesis.

### Preparation of whole arrested embryos

Embryos that showed no evidence of further development by either cell count or compaction and failed to develop to the expanded blastocyst stage by day 7 or showing signs of degeneration were considered arrested in development if no change was seen by time-lapse analysis for at least the preceding 24 hours. Arrested embryos were scored as either early, mid, or late as follows: early arrest, 1 cell through 6-10 cells to cover post-fertilization through embryonic genome activation; mid arrest, >10 cells to pre-compaction; and late arrest, evidence of compaction through early blastocoel formation. The zona pellucida of each arrested embryo was first thinned by brief exposure to acidified Tyrode’s solution (Origio, USA) and removed by gentle pipetting, ensuring the integrity of the entire intact cell mass for subsequent genetic analysis. All selected arrested embryos were then prepared for genetic analysis. DNA isolation, sample preparation, and whole genome amplification (WGA) was carried out as previously described (Gorodeckaja *et al.*, 2019).

### Data processing, statistical analysis, and visualization

Raw sequencing data were processed and visualized using Bluefuse Multi software (Illumina Inc, CA). A total of 30 samples (3.2%; 18 arrested embryos and 12 TE biopsies) were excluded from downstream analysis due to failed quality control (lack of DNA, excessive technical noise, or evidence of contamination based on negative controls).

Intra-sample chromosome mosaicism is diagnostic of mitotic error and is expected to produce copy number results intermediate between those expected of uniform trisomies or monosomies (full copy number changes) and disomy. We previously validated the threshold for meiotic aneuploidy as 70% of full copy number changes by parallel analysis of polar bodies and embryo samples by SNP genotyping and karyomapping (Handyside *et al.*, 2021). To allow for technical variability, values below 30% were considered as putative normal disomies. Most of the non-meiotic, putative mitotic aneuploidies had copy numbers in the range of 30-70% although a minority exceeded the 70% threshold and may have resulted from chromosome malsegregation in the early mitotic cleavage divisions. While it is likely therefore that a small proportion of embryos were mis-classified with this approach, we emphasize that the same thresholds were applied across all samples, thereby ensuring robustness of comparisons between embryos at different stages or with different developmental outcomes. We also acknowledge that precise copy number displacements are less reliable when multiple chromosomes are simultaneously affected by aneuploidy, though this limitation again equally affects all categories of embryos. Examples of copy number plots are provided in Figure S7.

Sex chromosome aneuploidies and segmental aneuploidies were not stratified into meiotic and mitotic aneuploidy categories, because the probability of mis-classification is much higher for these groups. In the case of sex chromosome aneuploidies, the baseline copy number expectations depend on the assumed sex of the embryo, which itself must be inferred. In the case of segmental aneuploidies, the smaller affected regions result in a lower ratio of signal to noise in normalized read depth. Moreover, individual genomic bins may encompass CNV breakpoints, causing the erroneous detection of intermediate copy number segments.

All statistical analysis was conducted in R (version 4.1.2). Figures were produced using the ‘ggplot2’ (Wickham, 2011) and ‘ggsankey’ (https://github.com/davidsjoberg/ggsankey) packages.

The relationships between probability of embryo arrest and various predictor variables (Figure 4) were modeled using binomial generalized linear mixed effects models (GLMMs), implemented with the ‘GLMMadaptive’ package (https://drizopoulos.github.io/GLMMadaptive/). Patient identifier was included as a random effect in all models, thereby addressing the non-independence among sets of embryos sampled from the same patient. Specifically, in the first model, the response variable was a binary indicator denoting whether or not the embryo arrested, while the copy number result category was the sole categorical predictor variable. The second model used the same response variable, but the total number of whole aneuploid autosomes (meiotic or mitotic) was the sole numeric predictor variable. The third model again used the same response variable, but the number of cells after the first mitotic division was included as the sole categorical predictor variable, where the possible categories were 1, 2, 3, 4, and >4. In all cases, coefficient estimates and 95% confidence intervals were converted from the logit to probability scale to facilitate interpretation. As coefficient estimates from GLMMs are conditional on the level of the random effect (i.e., patient), we report the average marginal effect (AME), which averages over all levels of the random effect.

The relationship between aneuploidy and day of blastocyst biopsy was modeled using a linear mixed effects model (LMM), implemented with the ‘lme4’ package (Bates et al., 2015), with AMEs computed using the ‘margins’ package (https://thomasleeper.com/margins/).

Key results were replicated upon restricting analysis to the subset of IVF cases where all embryos were tested with PGT-A (see Supplementary Materials).

## Data and code availability

De-identified chromosome-level aneuploidy calls and time lapse results for all tested embryos are available on Zenodo (doi: 10.5281/zenodo.6789485), along with deidentified metadata for the corresponding patients and cycles. Code necessary for reproducing all analyses and figures is available at https://github.com/mccoy-lab/embryo arrest and archived on Zenodo (doi: 10.5281/zenodo.6789485).

## Acknowledgments

The authors wish to thank the nursing team at the Bridge Clinic, London Women’s Clinic and clinical embryology team at London Women’s Clinic, Harley Street, London, for their assistance during this work. Thank you to the Origins of Aneuploidy Research consortium for facilitating this collaboration. Thank you also to Sara Carioscia for constructive comments on the manuscript. RCM is supported by the National Institute of General Medical Sciences of the National Institutes of Health under Award R35GM133747. The content is solely the responsibility of the authors and does not necessarily represent the official views of the National Institutes of Health.

## Declaration of interests

RCM is co-inventor on a provisional patent application by Johns Hopkins University related to inferring the origins of aneuploidies from PGT-A data. AHH, MCS, CO, AM and KA declare no competing interests.

## Notes

https://doi.org/10.5281/zenodo.6789485

### Summary of Updates

The revision includes edits to:provide more context to motivate our study in relation to previous literature investigating the links between chromosome abnormalities and embryo arrest; emphasize the novelty of our work in testing embryos regardless of their survival or morphology, thereby overcoming common biases associated with retrospective analyses of embryos that are candidates for IVF transfer; cite our validation study and related literature, providing supporting analyses in the Supplementary Materials, and more transparently discussing the strengths and limitations of the sequencing-based PGT-A platform.

