## Supplementary Materials for "Meiotic and mitotic aneuploidies drive arrest of in vitro fertilized human preimplantation embryos"

### Supplementary Figures

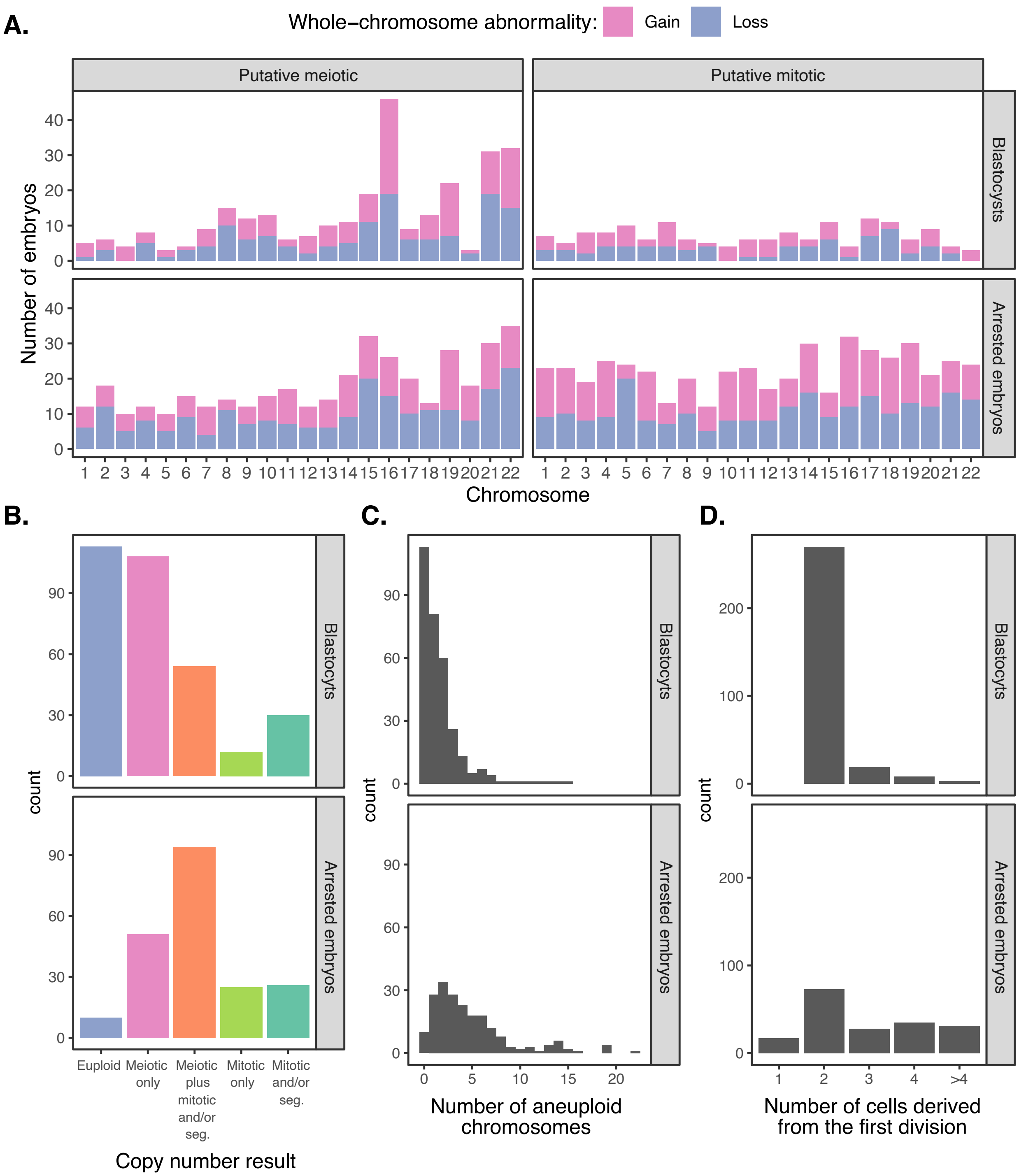


**Figure S1.** Characteristics of blastocysts and arrested embryos (see Fig. 2) but restricted to IVF cases where all blastocysts and arrested embryos were tested. **A.** Chromosome-specific counts of putative meiotic (i.e., full copy number change) and putative mitotic (i.e., intermediate copy number change) whole-chromosome gains and losses observed in blastocysts and arrested embryos as determined with PGT-A. Only autosomes are depicted, as distinguishing meiotic and mitotic origins of aneuploidies affecting sex chromosomes poses unique challenges (see Methods). **B.** Counts of arrested embryos versus developing blastocysts, stratified by PGT-A copy number result category. **C.** Counts of arrested embryos versus developing blastocysts, stratified by total number of aneuploid chromosomes. **D.** Counts of arrested embryos versus developing blastocysts, stratified by number of cells present after the first mitotic division.

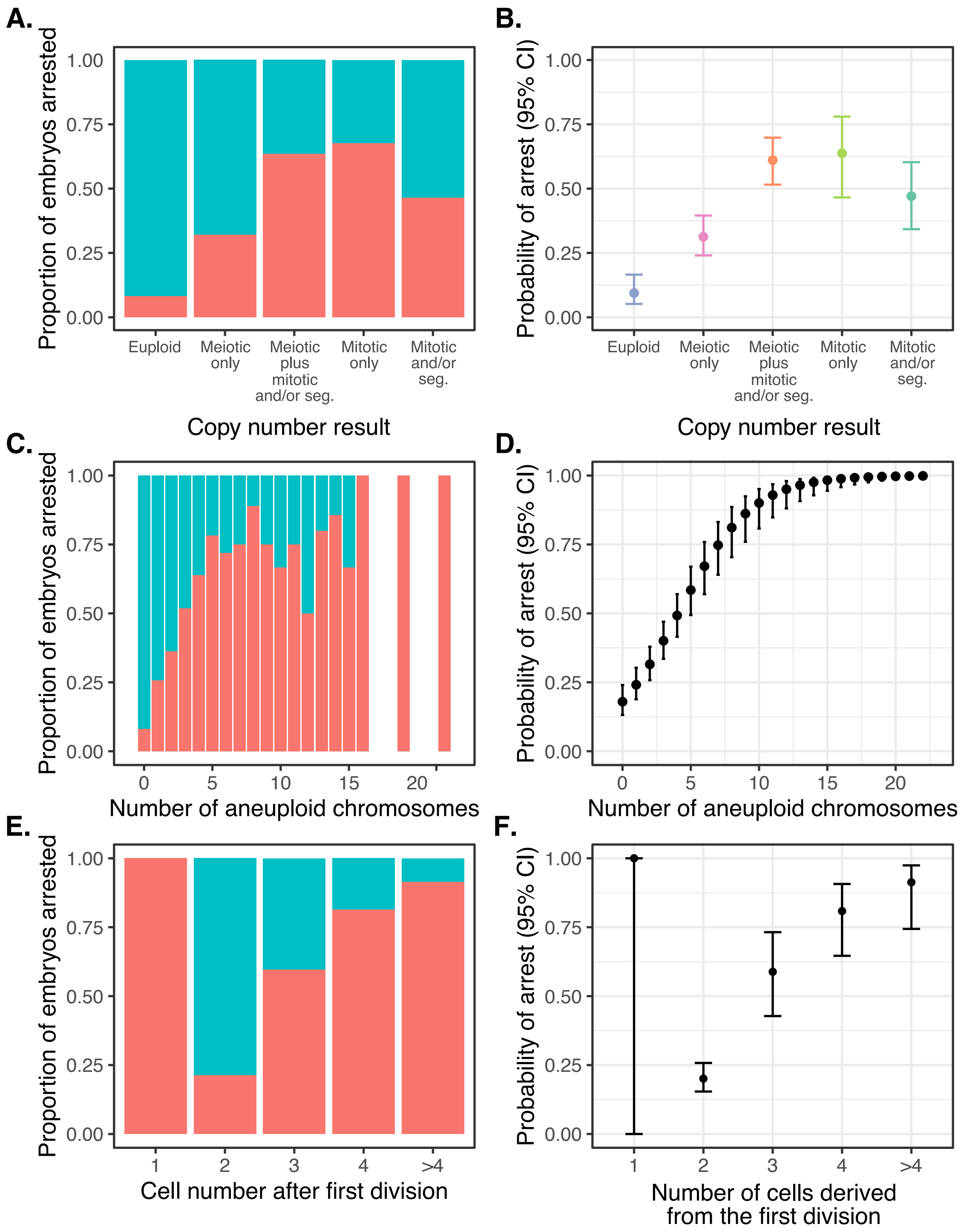


**Figure S2.** Data (left panels) and statistical modeling (right panels) of the proportion/probability of embryo arrest, stratifying on various patterns of chromosome copy number or cell division. Analyses are identical to those presented in Fig. 3 but restricted to IVF cases where all blastocysts and arrested embryos were tested. Error bars denote 95% confidence intervals of estimates. **A.** Proportion of embryos arrested (red) versus unarrested (blue), stratifying on chromosome copy number pattern, as assessed by PGT-A. **B.** Statistical modeling of the data from panel A. **C.** Proportion of embryos arrested (red) versus unarrested (blue), stratifying on the number of aneuploid chromosomes. **D.** Statistical modeling of data from panel C. **E**. Proportion of embryos arrested (red) versus unarrested (blue), stratifying on the number of cells observed after the first mitotic division, where 2 is normal. **F.** Statistical modeling of data from panel E.

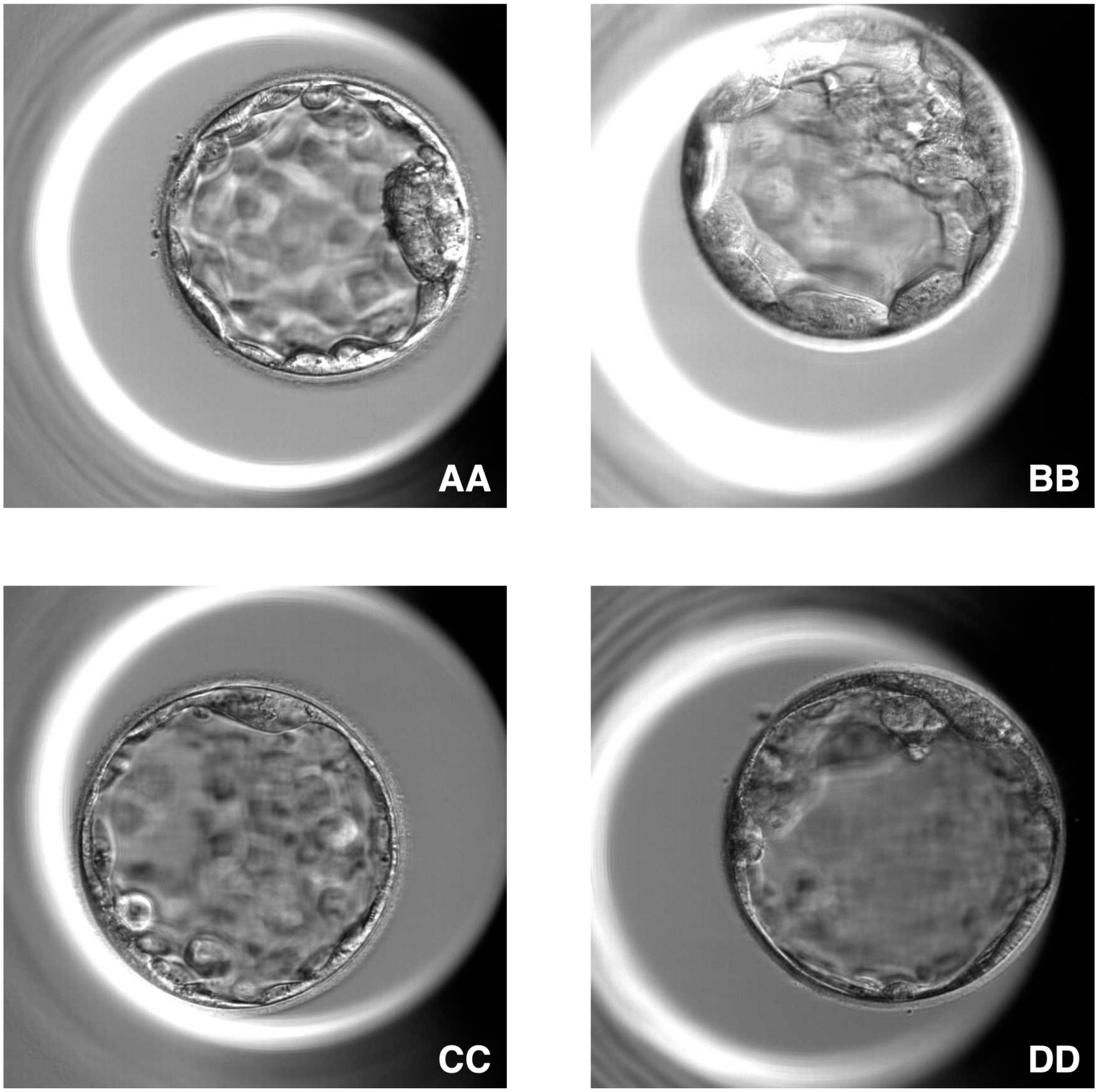


**Figure S3.** Examples of embryos of various grades as determined according to Alpha/ESHRE (2011) guidelines. Embryos with grades AA, BB, CC, and DD are depicted.

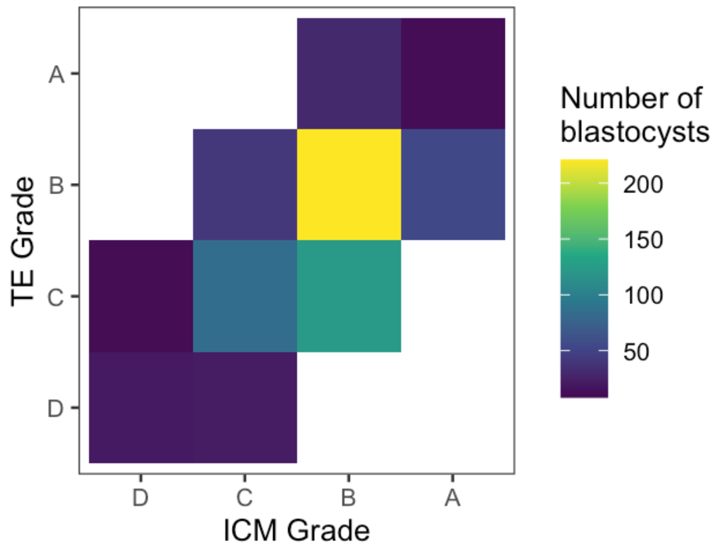


**Figure S4.** Numbers of embryos with various combinations of TE and ICM morphological grades.

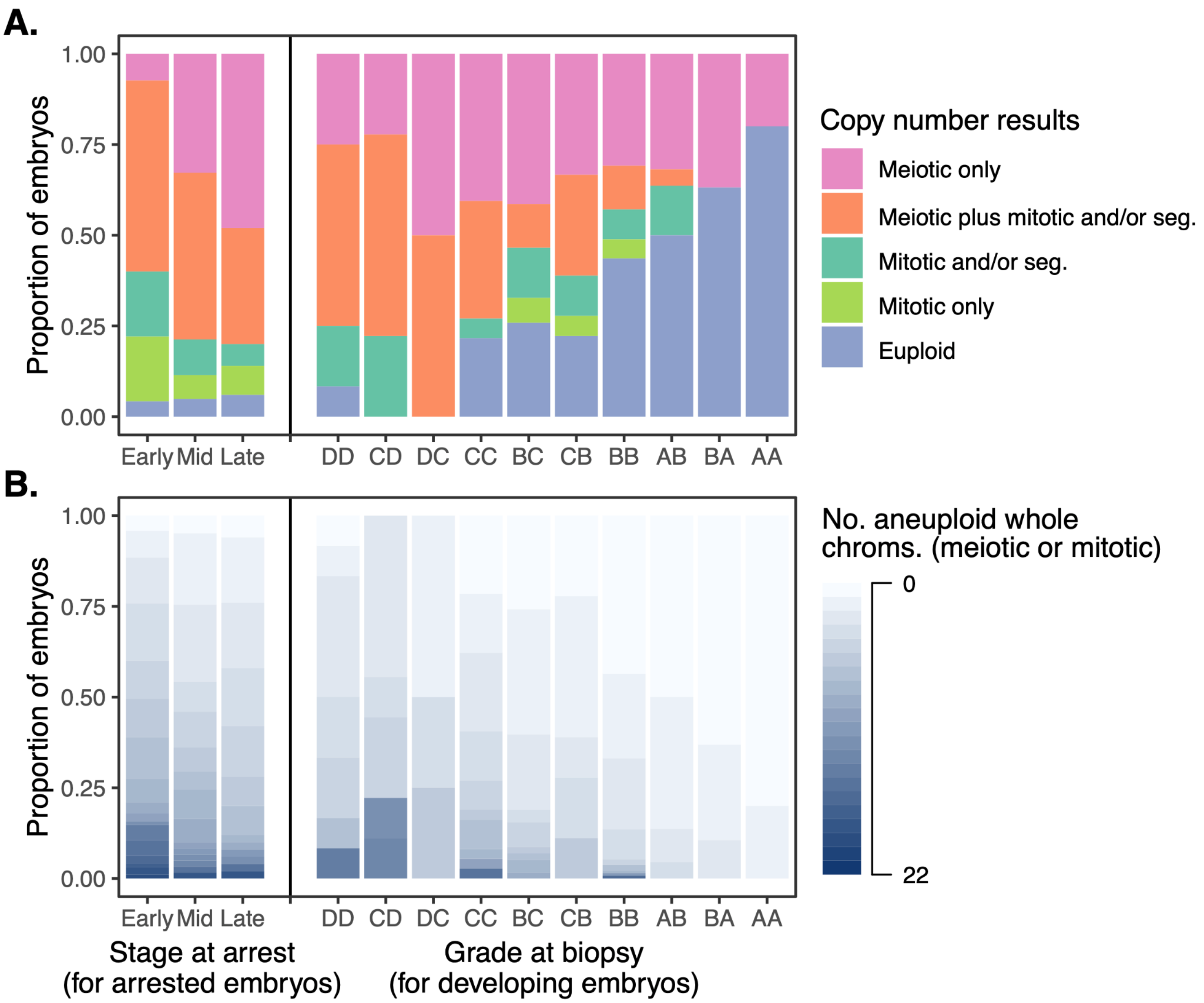


**Figure S5.** Chromosome copy number results (as assessed via PGT-A) across all tested embryos, stratifying by stage at arrest (see Methods for description of arrested embryos) or morphological grade (for embryos that formed blastocysts). Analyses are identical to those presented in Fig. 4 but restricted to IVF cases where all blastocysts and arrested embryos were tested. ICM grade is listed first, and TE grade is listed second. **A.** Copy number results assigned to categories, as described in Table 1. **B.** Copy number results depicted as counts of aneuploid chromosomes.

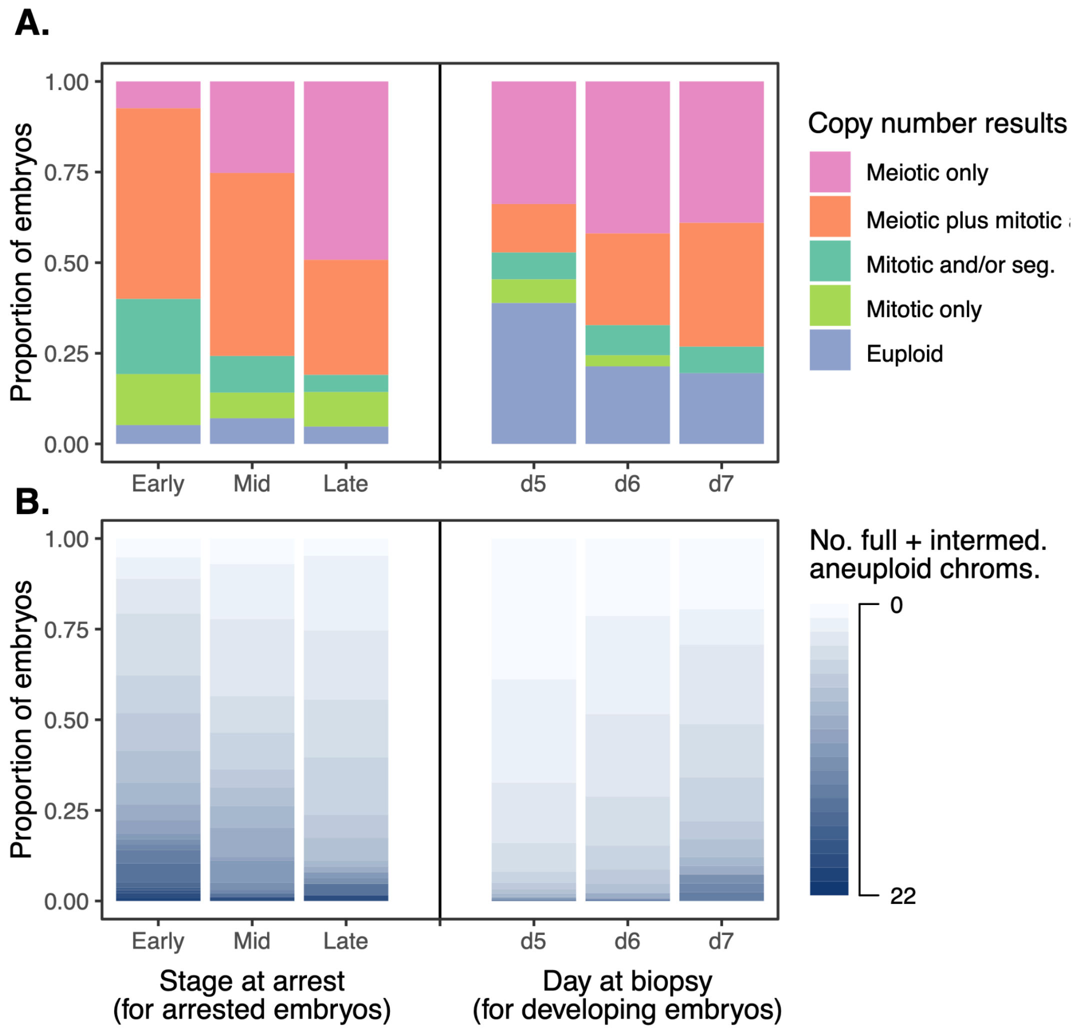


**Figure S6.** Chromosome copy number results (as assessed via PGT-A) across all tested embryos, stratifying by stage at arrest (see Methods) or day of biopsy (for embryos that formed blastocysts). **A.** Copy number results assigned to categories, as described in Table 1. **B.** Copy number results depicted as counts of aneuploid chromosomes.

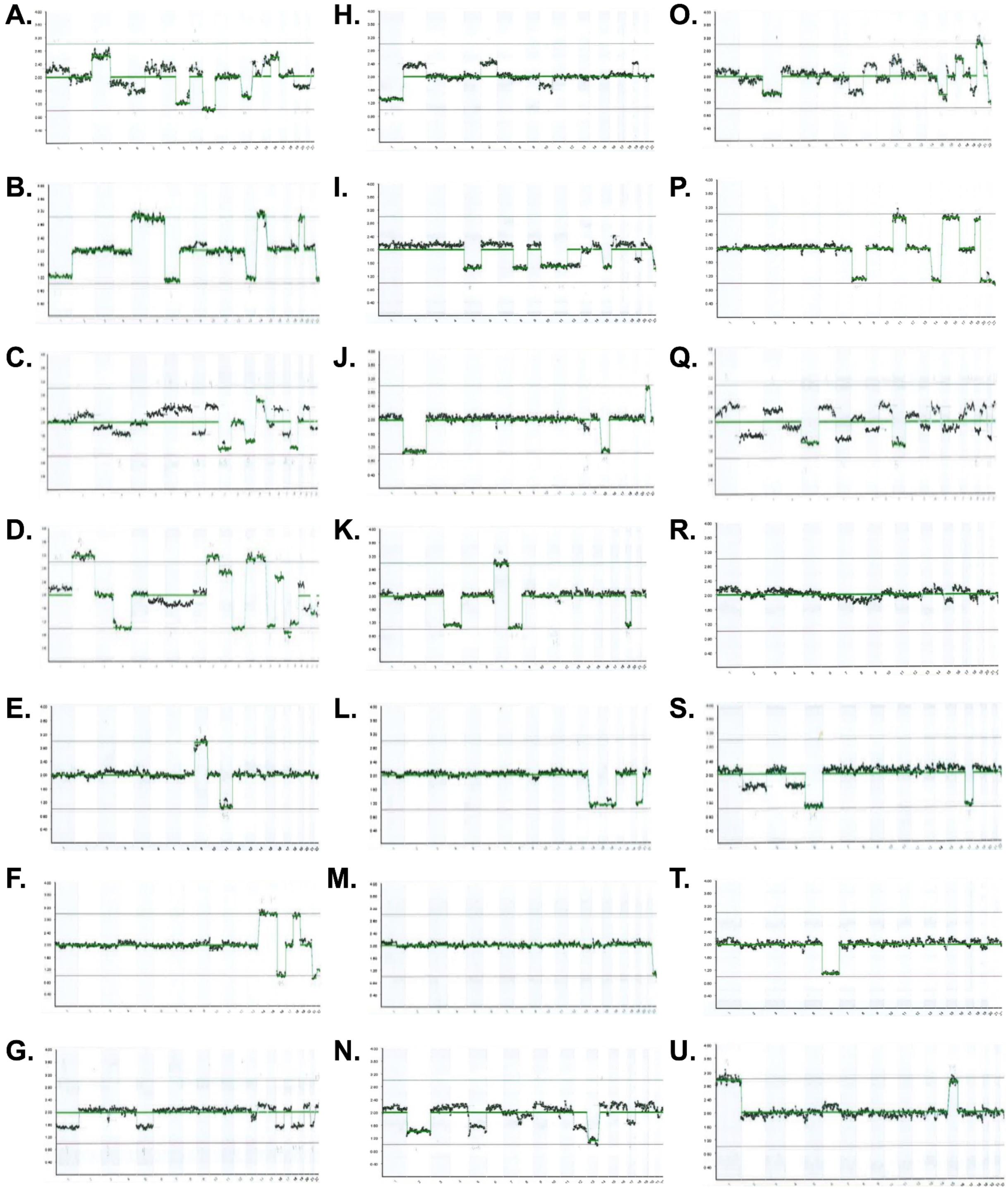


**Figure S7.** Representative examples of PGT-A results from which chromosome copy number measurements were obtained. Chromosomes are arranged along the X-axes, while inferred copy numbers are depicted on the Y-axes. Upper and lower bounding lines (gray) indicate expectations for full (i.e., putative meiotic) trisomy and monosomy, respectively, while the mid-line indicates expectations for diploidy. Corresponding cell division patterns and survival status are indicated as follows. **A.** Multipolar first division; 1-4-8 cells; mid arrest. **B.** Normal division; 1-2-4 cells; mid arrest. **C.** Tripolar first division; 1-3-6 cells; early arrest. **D.** Abnormal first division; 1-3-6 cells; early arrest. **E.** No division; 1 cell; early arrest. **F.** Abnormal first division; 1-3-6 cells; mid arrest. **G.** Abnormal first division; 1-4-8 cells; mid arrest. **H.** Tripolar first division; precocious second division; 1-4 cells; early arrest. **I.** Tripolar first division; 1-3-6 cells; mid arrest. **J.** Abnormal second division; 1-2-7 cells; mid arrest. **K.** Abnormal first division; 1-3-10 cells; late arrest. **L.** Abnormal first division; 1-3-9 cells; mid arrest. **M.** Normal division; 1-2-4 cells; late arrest. **N.** Tripolar first division; 1-3-6 cells; late arrest (with cells excluded at compaction). **O.** Abnormal first division; 1-4 cells; mid arrest. **P.** Normal division; 1-2-4 cells; late arrest. **Q.** Abnormal first and second division; 1-4-7 cells; early arrest. **R.** Day-6 expanded blastocyst; 4DD. **S.** Day-6 expanded blastocyst; 4DD. **T.** Day-6 expanded blastocyst; 4DD. **U.** Day-7 expanded blastocyst; 4DD.

#### Supplementary Tables

| | | Total counts | | | Counts per patient | | |
|---|---|---|---|---|---|---|---|
| **Maternal age** | **n** | **2PN** | **Bs** | **Arrested** | **2PN** | **Bs** | **Arrested** |
| **30** | 1 | 5 | 3 | 2 | 5.0 | 3.0 | 2.0 |
| **31** | 3 | 29 | 24 | 5 | 9.7 | 8.0 | 1.7 |
| **32** | 4 | 14 | 8 | 6 | 3.5 | 2.0 | 1.5 |
| **33** | 4 | 18 | 8 | 10 | 4.5 | 2.0 | 2.5 |
| **34** | 9 | 63 | 42 | 21 | 7.0 | 4.6 | 2.3 |
| **35** | 11 | 88 | 47 | 41 | 8.0 | 4.2 | 3.7 |
| **36** | 8 | 72 | 40 | 32 | 9.0 | 5.0 | 4.0 |
| **37** | 10 | 67 | 42 | 25 | 6.7 | 4.2 | 2.5 |
| **38** | 12 | 119 | 61 | 58 | 9.9 | 5.1 | 4.8 |
| **39** | 14 | 82 | 40 | 42 | 5.9 | 2.9 | 3.0 |
| **40** | 19 | 137 | 86 | 51 | 7.2 | 4.5 | 2.7 |
| **41** | 16 | 103 | 56 | 47 | 6.4 | 3.5 | 2.9 |
| **42** | 23 | 194 | 91 | 103 | 8.4 | 4.0 | 4.5 |
| **43** | 18 | 112 | 48 | 64 | 6.2 | 2.7 | 3.6 |
| **44** | 9 | 65 | 21 | 44 | 7.2 | 2.3 | 4.9 |
| **45** | 4 | 12 | 5 | 7 | 3.0 | 1.3 | 1.8 |
| **Total** | 165 | 1180 | 622 | 558 | 7.1 | 3.8 | 3.4 |

**Table S1.** Maternal age versus the number of dipronuclear (2PN) embryos developing to the blastocyst (Bs) stage. Data exclude 22 cycles where all embryos arrested and were not tested by PGT-A (a total of 52 2PN zygotes)

| Type of first division | n | Rate of aneuploidy | Estimated probability of arrest (95% CI) | Type of second division | n | Rate of aneuploidy | Estimated probability of arrest (95% CI) |
|---|---|---|---|---|---|---|---|
| **Normal** | 621 | 72.0% | 28.1%<br>(22.2% - 34.8%) | **Normal** | 540 | 70.2% | 22.7% (17.7% - 28.7%) |
| | | | | **Multipolar** | 40 | 82.5% | 69.6% (46.8% - 85.7%) |
| | | | | **Precocious** | 27 | 81.5% | 72.3% (48.4% - 87.9%) |
| | | | | **Failed** | 12 | 91.7% | 59.0% (27.7% - 84.5%) |
| | | | | **Reverse** | 2 | 100% | 100% (0% - 100%) |
| **Precocious** | 171 | 86.5% | 76.8%<br>(67.1% - 84.3%) | **Normal** | 53 | 81.1% | 45.5% (30.3% - 61.6%) |
| | | | | **Precocious** | 14 | 85.7% | 91.7% (54.4% - 99.0%) |
| | | | | **Failed** | 10 | 80% | 26.4% (5.9% - 67.3%) |
| | | | | **Multipolar** | 7 | 85.7% | 100% (0% - 100%) |
| | | | | **Reverse** | 1 | 100% | 100% (0% - 100%) |
| | | | | **Degenerate** | 1 | 100% | 100% (0% - 100%) |
| **Multipolar** | 29 | 93.1% | 84.3%<br>(61.3% - 94.8%) | **Normal** | 16 | 87.5% | 77.9% (46.4% - 93.5%) |
| | | | | **Precocious** | 2 | 100% | 100% (0% - 100%) |
| | | | | **Reverse** | 2 | 100% | 100% (0% - 100%) |
| | | | | **Multipolar** | 1 | 100% | 100% (0% - 100%) |
| | | | | **Failed** | 1 | 100% | 100% (0% - 100%) |
| **Failed** | 22 | 100% | 100%<br>(0% - 100%) | **Failed** | 8 | 100% | 100% (0% - 100%) |
| | | | | **Normal** | 7 | 100% | 100% (0% - 100%) |
| | | | | **Precocious** | 4 | 100% | 100% (0% - 100%) |
| | | | | **Multipolar** | 3 | 100% | 100% (0% - 100%) |

**Table S2.** Outcomes of first and second cleavage divisions as measured from time-lapse data, along with associated rates of aneuploidy (any form) and estimated probabilities of arrest. Note that in cases of an abnormal first division, the type of abnormal second division was not always possible to classify, so the total sample size for the second division is less than that of the first division. "Multipolar" cleavage refers to the direct cleavage of the zygote (or a daughter cell) into three or more cells. "Precocious" cleavage refers to a rapid division pattern where the zygote (or a daughter cell) undergoes a normal 1→2 cell cleavage, followed by a subsequent premature division to produce 3 or more blastomeres. "Reverse" cleavage refers to the resorption of blastomeres after cytokinesis. "Failed" cleavage refers to multiple rounds of karyokinesis without cytokinesis.

| ICM | TE |
|---|---|
| A : ICM prominent, easily seen, tightly adhered compacted cells | A : Continuous layer of small identical cells |
| B : ICM less prominent (cells appear compacted and larger in size, loosely adhered) | B : Fewer cells with gaps, not continuous |
| C : Very few cells visible (cells similar to TE) | C : Fewer small cells with large cells, not continuous |
| D : No visible cells or visible cells are degenerate or necrotic | D : Sparse cells, large/flat/degenerate |

**Table S3.** A description of the embryo grading scheme employed in our study, which was obtained from the ACE/NEQAS guidelines and is aligned with Alpha/ESHRE recommendations (Alpha/ESHRE, 2011).
